# Perceived threat bias and reduced hippocampal volume in combat veterans

**DOI:** 10.1101/313221

**Authors:** Daniel W. Grupe, Benjamin A. Hushek, Kaley Ellis, Andrew J. Schoen, Joseph Wielgosz, Jack B. Nitschke, Richard J. Davidson

**Author notes:** Corresponding author Daniel W. Grupe, Ph.D. Center for Healthy Minds, 625 West Washington Ave, Madison, WI 53703. **Disclosures/Conflicts:** Dr. Davidson is the founder and president, and serves on the board of directors, for the non-profit organization Healthy Minds Innovations, Inc. The other authors report no conflicts of interest or financial disclosures.

## Abstract

**Background:** Reduced hippocampal volume is frequently observed in posttraumatic stress disorder (PTSD), but the exact psychological processes associated with these alterations remain unclear. Given the role of the hippocampus in contextual representations of threat and memory, we investigated relationships between retrospectively reported combat exposure, perceived threat, and hippocampal volume in trauma-exposed veterans.

**Methods:** T1-weighted anatomical MRI scans were obtained from 52 male veterans with a broad range of PTSD symptoms. Hippocampal volume was estimated using automatic segmentation tools in FreeSurfer. An index of perceived threat bias was calculated, reflecting the degree of discordance between subjective perceptions of threat while deployed and self-reported combat exposure. Hippocampal volume was regressed on perceived threat bias and PTSD symptoms on the Clinician-Administered PTSD Scale (CAPS).

**Results:** Perceived threat bias was unrelated to overall CAPS symptoms, but was positively correlated with CAPS avoidance/numbing symptoms and symptoms of depression, anxiety, and worry. The degree of perceived threat bias was inversely correlated with hippocampal volume. Hippocampal volume was also inversely related to avoidance/numbing CAPS symptoms.

**Conclusions:** These results indicate that volume of the hippocampus, a region involved in contextual threat processing and memory, is related to recalled associations between traumatic events and accompanying subjective threat appraisals. Future research should clarify the precise temporal milieu of these effects and investigate whether individual differences in hippocampal structure and function contribute to exaggerated threat appraisal at the time of trauma, or in subsequently biased memories or appraisals of traumatic events.

## Introduction

Reduced hippocampal volume is consistently observed in posttraumatic stress disorder (PTSD), with meta-analyses revealing these reductions across different trauma types and demographic groups (Karl et al., 2006; Kühn and Gallinat, 2013; Woon et al., 2010). The magnitude of this reduction, however, is quite modest: the largest study of subcortical structures in PTSD – a retrospective multi-site study consisting of nearly 1,900 participants – revealed an effect size of *d* = 0.17 for participants with PTSD vs. trauma-exposed controls (Logue et al., 2018). Exposure to trauma alone, even in the absence of PTSD pathology, can be associated with volumetric reductions, as hippocampal volume in trauma-exposed controls falls between that of individuals with PTSD and unexposed controls (Karl et al., 2006; Woon et al., 2010). These findings of modest effect sizes and morphometric differences in the absence of PTSD symptoms highlight the need for updated models of what is reflected in post-traumatic structural alterations to the hippocampus.

A separate line of research has investigated psychological and psychosocial risk factors to explain why different individuals exposed to similar traumatic events experience divergent long-term trajectories (King et al., 2006). This question, however, may be based on a faulty premise: just because two individuals are exposed to the same external circumstances does not mean that they experienced the “same” trauma. The interpretation and meaning of these traumatic experiences will vary widely across individuals based on biological and psychological predispositions, past experiences, and current environmental factors. Indeed, subjectively *perceived threat* – fear or worry about one’s safety and well-being during and after exposure to trauma – is one of the best predictors of PTSD (Blanchard et al., 1995; King et al., 2006; Lancaster et al., 2016) and other mood and anxiety disorders (King et al., 2008; Mott et al., 2012), and mediates the relationship between combat exposure and PTSD symptoms in multiple veteran samples (King et al., 1995; Vogt et al., 2011; Vogt and Tanner, 2007). These relationships between subjective threat appraisals and the emergence of psychopathology underscore the importance of identifying neurobiological mechanisms of this psychological characteristic.

The hippocampus is a prime candidate region that may be related to subjectively perceived threat, due to its central role in the contextual processing of threat (Liberzon and Abelson, 2016) and the aforementioned evidence for structural alterations to the hippocampus following trauma exposure. Two studies of healthy, older adults (Gianaros et al., 2007; Zimmerman et al., 2016) identified an inverse relationship between hippocampal volume and the related construct of perceived stress, the degree to which individuals appraise daily life events as being stressful, overwhelming, and uncontrollable. However, no studies to our knowledge have directly examined the relationship between perceived threat following combat trauma and hippocampus structure.

An important consideration in studying perceived threat in combat-exposed individuals is that deployment environments can be associated with objectively high levels of threat, in which case extreme levels of perceived threat would reflect contextually appropriate, adaptive responses. Whereas increased attentional biases toward threat in new military recruits predict the eventual development of PTSD during a safe baseline period, the opposite is true immediately prior to and during deployment, when increased threat avoidance is associated with later PTSD (Wald et al., 2013, 2011). It is not the case that particular behavioral profiles are universally adaptive or maladaptive; rather, a defining characteristic of maladaptive threat responding is incongruence between a specific context/environment and one’s response (Shechner and Bar-Haim, 2016). Examining perceived threat independent of the degree of trauma exposure may fail to distinguish adaptive from pathological modes of threat processing. In considering risk for psychopathology, the critical factor may not be absolute levels of perceived threat, but *relative biases* in perceived threat for individuals exposed to similar levels of traumatic events.

To that end, we developed a novel measure of “perceived threat bias”, which quantifies the relative incongruence of retrospectively reported combat exposure and perceived threat during deployment. In 52 combat-exposed veterans, we investigated relationships between perceived threat bias and symptoms of PTSD, depression, anxiety, and trait worry. We tested the hypothesis that elevated perceived threat bias would be associated with reductions in hippocampal volume, and also investigated hippocampal volume as a function of PTSD symptom severity. To investigate the specificity of relationships to the hippocampus, analogous analyses were conducted for the amygdala, due to its role in threat perception (van Wingen et al., 2011) and observations of smaller amygdala volume in PTSD (Karl et al., 2006; Morey et al., 2012).

## Methods

### Participants

We recruited veterans of Operation Enduring Freedom/Operation Iraqi Freedom through community and online advertisements and in collaboration with veterans’ organizations, the Wisconsin National Guard, and the Madison VA. Following complete study description, written informed consent was obtained. A team of clinically trained interviewers administered the Clinician-Administered PTSD Scale (CAPS; Blake et al., 1990) and Structured Clinical Interview for DSM-IV (SCID; First et al., 2002) with supervision from a licensed clinical psychologist (JBN). Exclusionary conditions included substance dependence within the past 3 months and lifetime bipolar, psychotic, or cognitive disorders.

Participants were enrolled either into a combat-exposed control (CEC) group or a posttraumatic stress symptoms (PTSS) group (Table 1). Participants in the CEC group had no current Axis I disorder and very low PTSD symptoms (CAPS scores < 10) and did not meet diagnostic criteria for any CAPS subscales. Participants in the PTSS group had PTSD symptoms occurring at least monthly with moderate intensity and CAPS scores ≥ 20 and met diagnostic criteria for at least 1 of 3 CAPS subscales. Current treatment with psychotropic medications (other than benzodiazepines or beta-blockers) or maintenance psychotherapy was permitted if treatment was stable for 8 weeks.

**Table 1:**
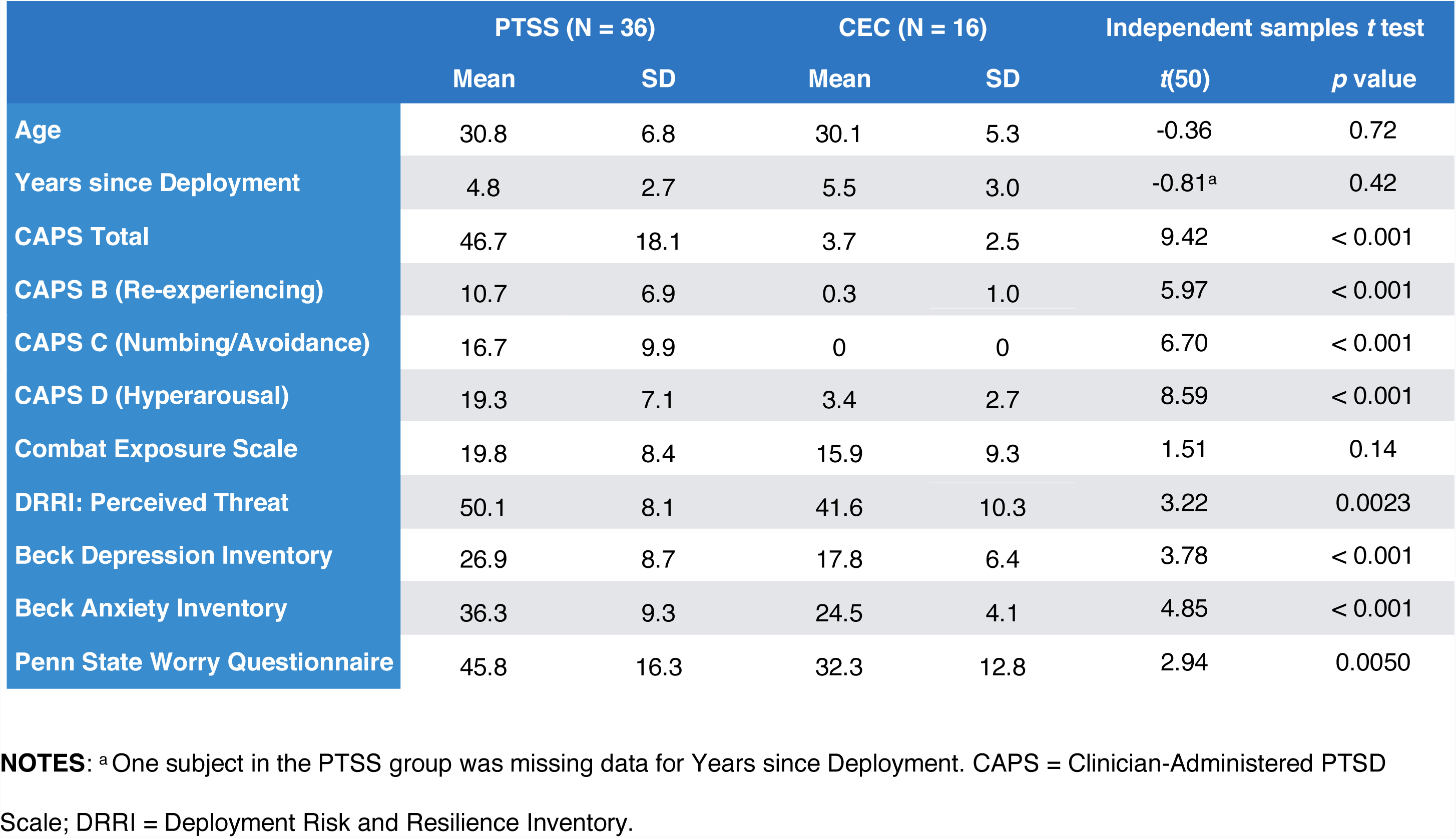
Demographic, clinical, and self-report data for the posttraumatic stress symptoms (PTSS) and combat-exposed control (CEC) groups.

58 veterans met eligibility criteria and were enrolled, but due to the small number of female veterans (n=4) we analyzed data from men only. Data from 2 participants were excluded due to excessive motion that prevented accurate delineation of white/gray matter boundaries. The final sample consisted of 36 PTSS subjects – 17 of whom met full PTSD diagnostic criteria and 19 who met diagnostic criteria for 1 or 2 of the CAPS subscales – and 16 veterans enrolled in the CEC group. We previously reported on relationships between PTSD symptoms and fMRI activation in an overlapping sample (Grupe et al., 2016).

### Data collection

In a pre-MRI visit, participants completed self-report measures including the Combat Exposure Scale (CES; Keane et al., 1989) subscales from the Deployment Risk and Resiliency Inventory (DRRI; King et al., 2006), the Beck Anxiety Inventory (BAI; Beck et al., 1988), Beck Depression Inventory (BDI-II; Beck et al., 1996), and Penn State Worry Questionnaire (Meyer et al., 1990). Participants took part in an MRI scan within the subsequent 40 days. MRI data were collected on a 3T X750 GE Discovery scanner using an 8-channel head coil and ASSET parallel imaging with an acceleration factor of 2. Brain structure was assessed through the collection of T1-weighted anatomical images with 1-mm isotropic voxels (“BRAVO” sequence, TR=8.16, TE=3.18, flip angle=12°, field of view=256 mm, 256x256 matrix, 156 axial slices). Self-reported PTSD symptoms were assessed on the day of the MRI scan using the PTSD Checklist-Military Version (PCL-M; Blanchard et al., 1996).

### Perceived threat bias calculation

The CES is a 7-item Likert scale assessing the frequency of different wartime stressors (example items: “Did you ever go on combat patrols or have other dangerous duty?”, “Were you ever under enemy fire?”). The DRRI includes 17 scales characterizing environmental, psychosocial, and psychological factors before, during, and after deployment. Among these scales is “perceived threat” (or “combat concerns”), a 15-item Likert scale reflecting veterans’ cognitive or subjective appraisals of combat-related danger (example items: “I thought I would never survive”, “I was concerned that my unit would be attacked by the enemy”).

We used these two scales to create an index of perceived threat bias (Figure 1). Both scales were approximately normally distributed across the entire sample (Figure 1A). To calculate perceived threat bias, we z-normalized each measure and calculated the difference between normalized perceived threat and normalized combat exposure scores (z(perceived threat) – z(combat exposure)). A perceived threat bias score of 1.0 thus reflects a 1 SD increase in perceived threat based on a given level of combat experience (Figure 1B).

**Figure 1:**
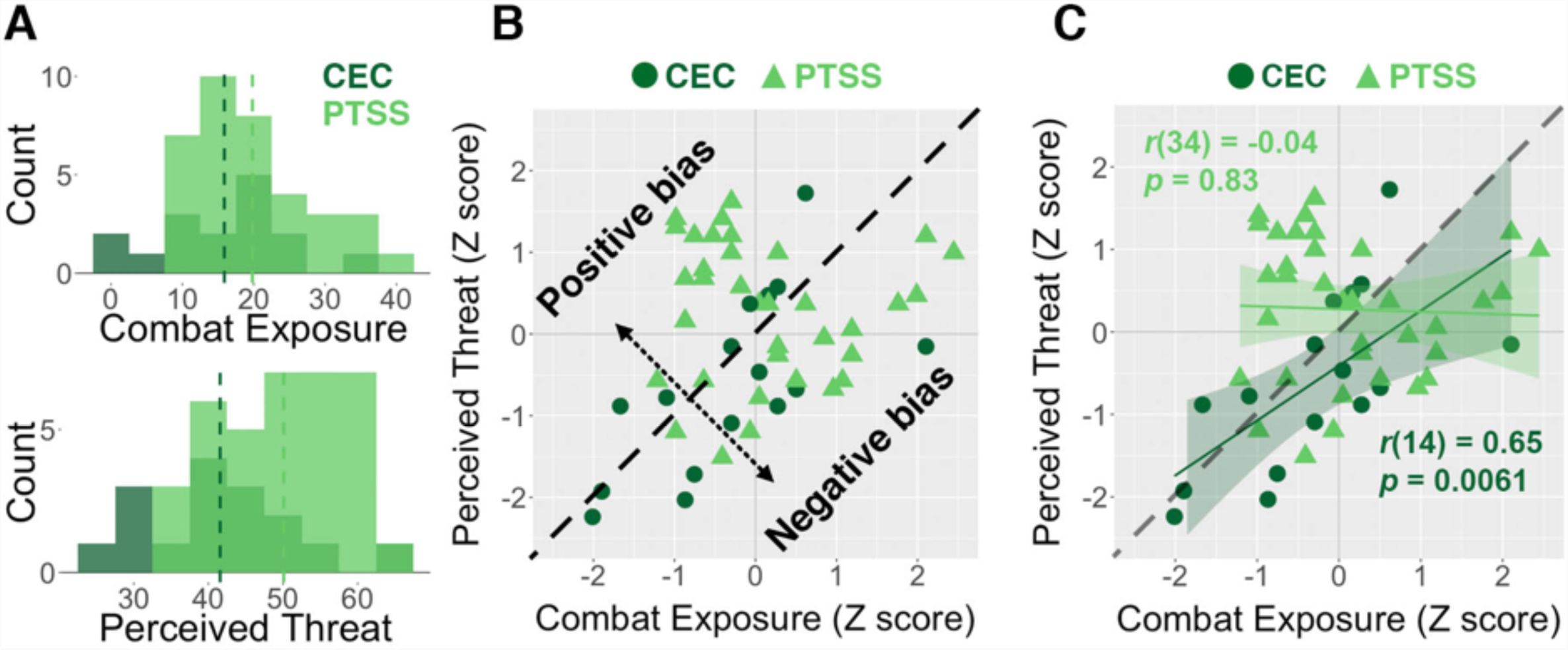
(A) Overlapping histograms show the distributions of self-reported combat exposure and perceived threat for the posttraumatic stress symptoms (PTSS) and combat-exposed control (CEC) groups. Each of these measures was approximately normally distributed across the entire sample, although the PTSS group had elevated perceived threat relative to the CEC group (*t*(50) = 3.22, *p* = 0.0023). Dashed lines indicate mean values for each group. (B) A plot of combat exposure vs. perceived threat, with the dashed line indicating zero perceived threat bias. Individuals above this line had a positive perceived threat bias, with elevated perceived threat relative to the amount of self-reported combat experience, whereas individuals below the line had a negative perceived threat bias. (C) Individuals in the CEC group showed a strong and positive correlation between combat experience and perceived threat, whereas this relationship was absent for individuals in the PTSS group. Shaded areas indicate 95% confidence intervals.

### FreeSurfer processing

Cortical reconstruction and volumetric segmentation were performed using the FreeSurfer image analysis suite (stable release version 5.3.0; http://surfer.nmr.mgh.harvard.edu/). Processing included motion correction, skull removal, intensity normalization, registration, segmentation of subcortical white and deep gray matter structures, white matter and pial surface tessellation, and cortical surface parcellation. Segmentation quality was visually assessed and manually edited as necessary (http://freesurfer.net/fswiki/Edits). Automated segmentation of the bilateral hippocampus and amygdala was conducted for each subject, a procedure that compares favorably with labor-intensive manual segmentation (Morey et al., 2009).

### Data analysis

To assess the convergent and discriminant validity of perceived threat bias in relation to existing measures, we calculated Pearson correlation coefficients with scores on the CAPS, BAI, BDI, and PSWQ across the entire sample and within the PTSS group. For the CAPS we examined correlations with total scores and each subscale. For each measure we also calculated correlations with CES and DRRI scores separately, allowing us to test whether one component was driving relationships observed for perceived threat bias, and whether this new construct provided incremental validity beyond existing scales of perceived threat and combat exposure.

We calculated correlations between total hippocampal and amygdalar volume and perceived threat bias scores to explore the neuroanatomical correlates of this novel measure. Correlations were conducted across the entire sample and within the PTSS group alone. We again calculated correlations between structural volume and CES/DRRI scores separately.

We correlated hippocampal and amygdalar volume with total CAPS symptoms in an effort to replicate previously observed volumetric reductions, and with each CAPS subscale to explore relationships with specific symptom clusters. We also compared hippocampal volume between the CEC group (N=16) and participants with a PTSD diagnosis (N=17).

Finally, we conducted exploratory voxelwise brain morphometry analyses to relate perceived threat bias and CAPS symptoms to structural changes within the anatomically constrained hippocampus and amygdala, and across the whole brain (Supplementary Materials).

## Results

### Clinical and self-report measures

Demographics and symptom data are provided in Table 1. The PTSS group had higher scores on the CAPS, BAI, BDI, and PSWQ. The groups did not differ on self-reported combat exposure on the CES, but the PTSS group did report elevated DRRI perceived threat.

### Perceived threat bias and relationships with symptoms/self-report data

Perceived threat bias was operationalized as the relative difference between self-reported combat exposure and perceived threat while deployed (see Methods). Consistent with previous research (Mott et al., 2012; Vogt et al., 2011), the CEC group showed a robust linear relationship between combat exposure and perceived threat (*r*(14) = 0.65, *p* = 0.0061). This pattern was absent in the PTSS group (*r*(34) = -0.04, *p* = 0.83; significant difference in correlation between groups, Fisher’s *Z* = 2.52, *p* = 0.011; Figure 1B).

There was no relationship between perceived threat bias and total CAPS scores in the full sample (*r*(50) = 0.16, *p* = 0.25) or in the PTSS group (*r*(34) = 0.15, *p* = 0.39; Figure 2A). However, perceived threat bias was correlated with emotional numbing/avoidance CAPS symptoms in the full sample (*r*(50) = 0.33, *p* = 0.01) and the PTSS group (*r*(34) = 0.29, *p* = 0.08; Figure 2B). Perceived threat bias was not associated with the other 2 PTSD symptom clusters, re-experiencing or hyperarousal (|*r*s| < 0.24, *p*s > 0.18; Table S1). Perceived threat bias was also correlated with self-reported avoidance/numbing symptoms on the PCL (full sample: *r*(50) = 0.39, *p* = 0.0045; PTSS group: *r*(34) = 0.32, *p* = 0.058) and, for the full sample only, total PCL symptoms (*r*(50) = 0.33, *p* = 0.019; PTSS group: *r*(34) = 0.04, *p* = 0.82).

**Figure 2:**
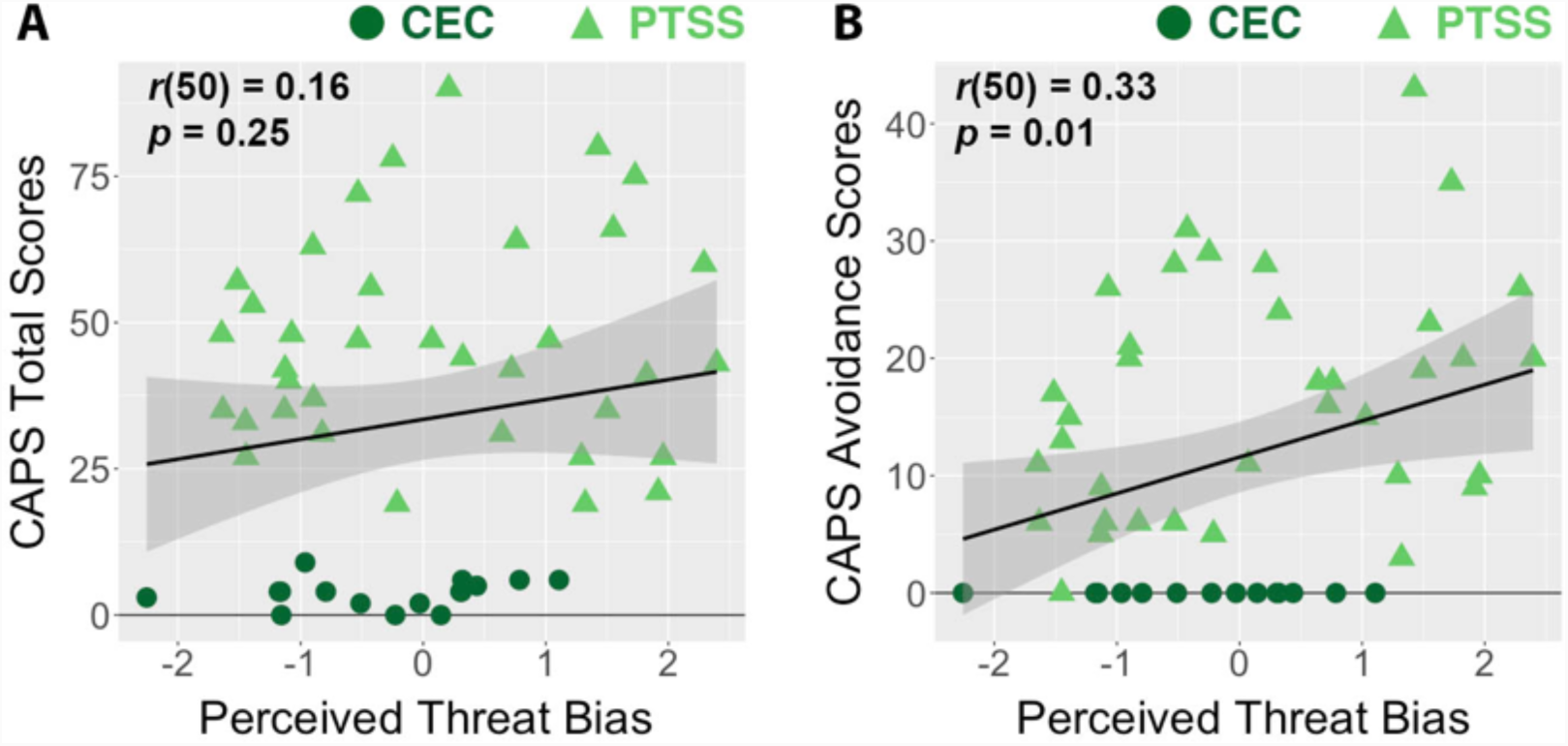
(A) Perceived threat bias showed no relationship with overall scores of PTSD on the Clinician-Administered PTSD Scale (CAPS) across the entire sample (*r*(50) = 0.16, *p* = 0.25) or in the posttraumatic stress symptoms (PTSS) group alone (*r*(34) = 0.04, *p* = 0.82). (B) There was a significant and positive correlation between perceived threat bias and CAPS avoidance/numbing symptoms across the entire sample (*r*(50) = 0.33, *p* = 0.01), and a non-significant correlation in the PTSS group (*r*(34) = 0.32, *p* = 0.06). Shaded areas indicate 95% confidence intervals.

Across the full sample and within the PTSS group, perceived threat bias was robustly correlated with depression symptoms on the BDI (*r*(50) = 0.48, *p* < 0.001), anxiety symptoms on the BAI (*r*(50) = 0.44, *p* = 0.001), and trait worry on the PSWQ (*r*(50) = 0.49, *p* < 0.001; Figure S1). In the full sample, these correlations were driven by individual differences in perceived threat, which showed similar correlations with symptom measures as did perceived threat bias (Table S2). In the PTSS group, however, symptoms of anxiety, depression, and worry were also *negatively* correlated with scores on the Combat Exposure Scale; as a result, correlations with these symptom measures in the PTSS group were numerically (but not statistically) higher for perceived threat bias vs. perceived threat alone (Table S2).

### Perceived threat bias is associated with smaller hippocampal volume

Hippocampal and amygdalar volumes for the entire sample and different subgroups of participants is provided in Table S3. Across all participants, elevated perceived threat bias was associated with reduced hippocampal volume (*r*(50) = -0.29, *p* = 0.039; Figure 3a). This relationship remained significant when adjusting hippocampal volume for total estimated intracranial volume (*r*(50) = -0.29, *p* = 0.035). Although the PTSS and CEC groups both showed inverse relationships between perceived threat bias and hippocampal volume (*r* = -0.25 and -0.41, respectively), a significant correlation was only seen when combining across all participants.

**Figure 3:**
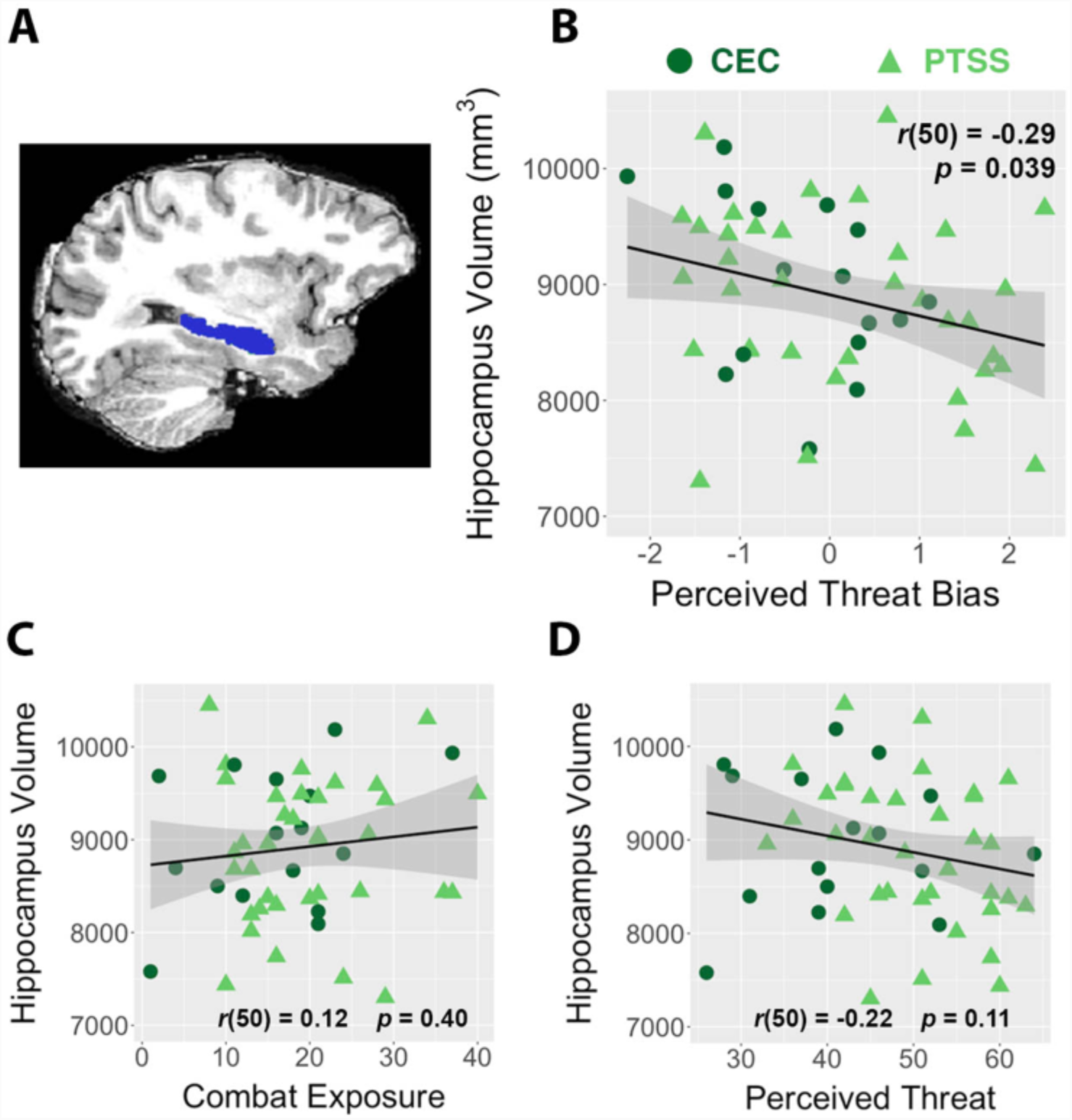
(A) Automatically segmented hippocampus ROI from a representative participant. (B) Across all participants, higher perceived threat bias scores were negatively correlated with bilateral hippocampal volume. Hippocampal volume was not significantly correlated with combat exposure (C) or perceived threat scores (D). Shaded areas indicate 95% confidence intervals.

We next assessed correlations between hippocampal volume and the subcomponents of perceived threat bias. Hippocampus volume was not significantly correlated with self-reported combat exposure (*r*(50) = 0.12, *p* = 0.40; Figure 3b) or perceived threat (*r*(50) = -0.22, *p* = 0.11; Figure 3c), although perceived threat bias and perceived threat did not differ statistically in their relationship with hippocampal volume (William’s *t*(50) = 0.57, *p* =0.57).

### PTSD avoidance/numbing symptoms are associated with smaller hippocampal volume

There was not a significant relationship between total PTSD symptoms on the CAPS and reduced hippocampal volume (*r*(50) = -0.21, *p* = 0.14; PTSS subjects: *r*(34) = -0.30, *p* = 0.079; Figure 4a). Similarly, there was no between-group difference in hippocampal volume between the CEC group and individuals diagnosed with PTSD (*t*(31) = 0.73, *p* = 0.47). We also investigated hippocampal volume in relation to specific CAPS symptom clusters. Hippocampal volume was most strongly related to avoidance/numbing symptoms (Figure 4c), with a non-significant relationship across all subjects (*r*(50) = -0.27, *p* = 0.052) and a significant relationship in the PTSS group (*r*(34) = -0.36, *p* = 0.032). Similar (and generally more robust) correlations with hippocampal volume were observed for PCL symptom scores (Figure S2).

**Figure 4:**
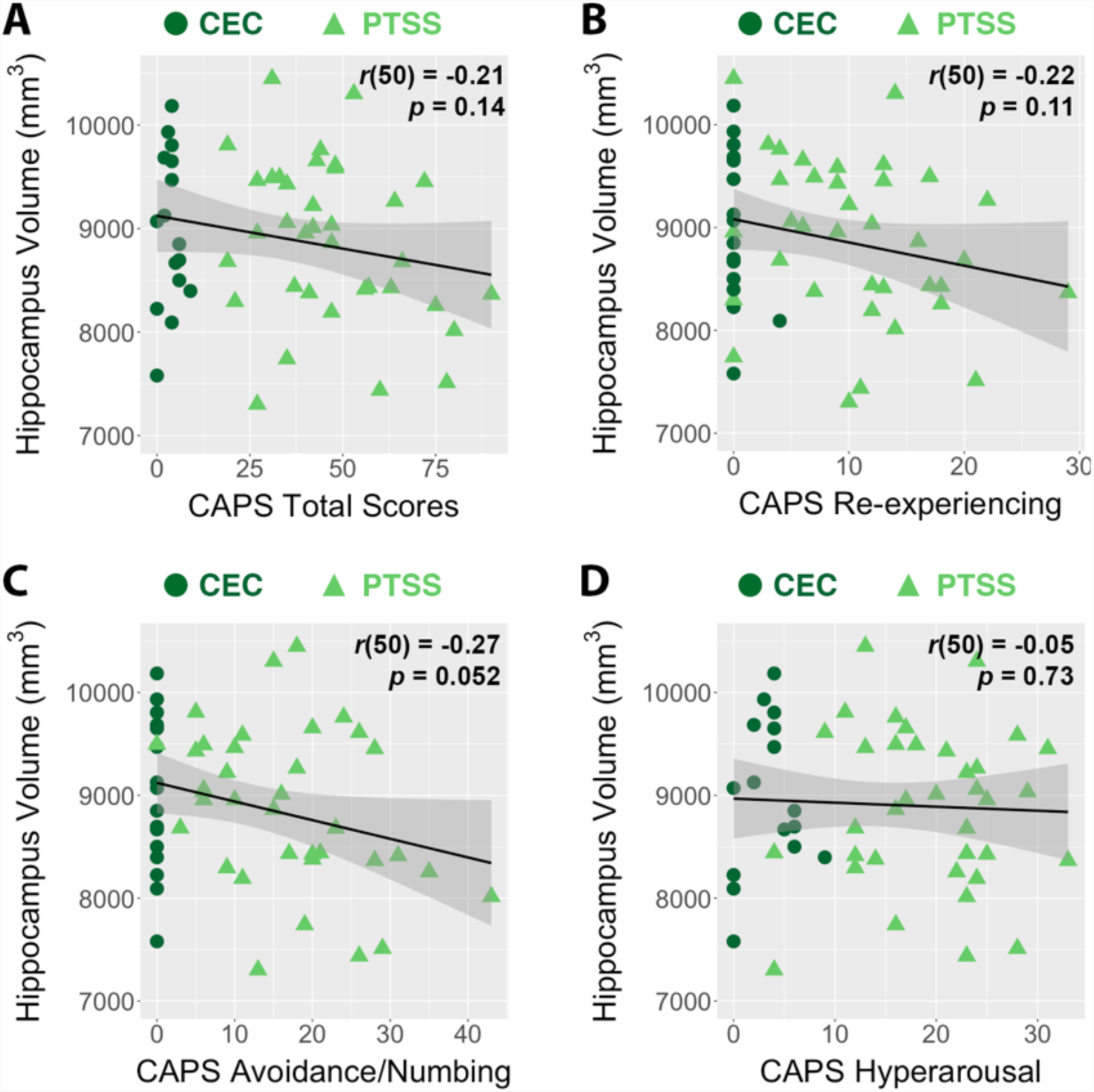
Overall PTSD symptom severity (A), as assessed by the Clinician-Administered PTSD Scale (CAPS), showed no significant relationship with bilateral hippocampal volume across the entire group (*r*(50) = -0.21, *p* = 0.14) or for subjects in the posttraumatic stress symptoms (PTSS) group alone (*r*(34) = -0.30, *p* = 0.079). Hippocampal volume was inversely correlated with CAPS avoidance/numbing symptoms (C) in the entire sample (*r*(50) = -0.27, *p* = 0.052) and the PTSS group alone (*r*(34) = -0.36, *p* = 0.032). Hippocampal volume was not related to re-experiencing (B) or hyperarousal symptoms (D) in the entire sample or the PTSS group (|*r*s| < 0.23, *p*s > 0.1).

Individual differences in perceived threat bias and CAPS avoidance/numbing symptoms were each associated with smaller hippocampal volume; further, perceived threat bias and avoidance/numbing symptoms were significantly correlated with one another. To see whether these factors explained shared or unique variance in hippocampal volume, we conducted a simultaneous linear regression of hippocampal volume on perceived threat bias and CAPS avoidance/numbing symptoms. In the full sample, neither perceived threat bias (*t*(49) = -1.57, *p* = 0.12) nor avoidance/numbing symptoms (*t*(49) = -1.39, *p* = 0.17) accounted for unique variance in hippocampal volume. The two factors together accounted for 8.2% of the variance in hippocampal volume (*F*(2,49) = 3.26, *p* = 0.047, multiple R^2^ = .118, adjusted R^2^ =.082). The analogous regression in the PTSS group showed that neither perceived threat bias (*t*(33) = -0.87, *p* = 0.39) nor avoidance/numbing symptoms (*t*(33) = -1.83, *p* = 0.076) accounted for unique variance in hippocampal volume. The two factors together accounted for 9.6% of the variance in hippocampal volume (*F*(2,33) = 2.85, *p* = 0.072, multiple R^2^ =.147, adjusted R^2^ = .096).

### Analysis of amygdalar volume in relation to perceived threat bias and PTSD symptoms

There were no relationships between amygdala volume and perceived threat bias (*r*(50) = 0.13, *p* = 0.35) or either perceived threat bias subcomponent (CES: *r*(50) = -0.17, *p* = 0.22; perceived threat: *r*(50) -0.02, *p* = 0.91). There were also no relationships between amygdala volume and total CAPS symptoms (*r*(50) = -0.10, *p* = 0.49) or individual CAPS symptom clusters (*r*s < 0.18, *p*s > 0.2), and there was no group difference in amygdala volume between the CEC group and individuals diagnosed with PTSD (*t*(31) = -0.15, *p* = 0.88).

### Whole-brain structural correlates of perceived threat bias and PTSD symptoms

In exploratory analyses we regressed voxelwise brain morphometry (VBM) maps on perceived threat bias and CAPS scores (Figure S3). Consistent with ROI-based volumetric analyses, at *p* < 0.1 (small volume correction within the amygdala/hippocampus), perceived threat bias was inversely correlated with local gray matter volume in the left mid/posterior hippocampus. This relationship was driven by perceived threat, which was significantly associated with reduced gray matter volume across much of the left hippocampus (*p* < 0.05). Combat Exposure Scale scores were *positively* associated with gray matter volume in the right posterior hippocampus at *p* < 0.1. CAPS scores were not associated with VBM in the hippocampus or amygdala, and there were no clusters that survived whole-brain correction for perceived threat bias, its subcomponents, or CAPS scores.

## Discussion

The relationship between combat experiences and maladaptive psychological outcomes has been shown to be mediated by subjective perceptions of threat (King et al., 2006; Vogt et al., 2011; Vogt and Tanner, 2007), but little is known about underlying neurobiological mechanisms. Here, we introduce the construct of perceived threat bias, reflecting relative discrepancies between self-reported combat experiences and perceived threat while deployed. In a sample of male OEF/OIF veterans, perceived threat and perceived threat bias were positively correlated with avoidance/numbing symptoms of PTSD, as well as depression and anxiety symptoms and trait worry. In addition, elevated perceived threat bias was associated with reduced hippocampal volume, suggesting that this brain region may play an important role in differential threat appraisals in the aftermath of combat exposure or other traumatic events.

Previous research on perceived threat has underscored the importance of this factor in conferring risk for trauma-related psychopathology (King et al., 2008; Mott et al., 2012; Vogt et al., 2011; Vogt and Tanner, 2007). Our measure of perceived threat bias acknowledges that the *mismatch* between trauma exposure and subjective threat appraisals may be a better indicator of maladaptive responding than absolute levels of perceived threat (Shechner and Bar-Haim, 2016; Wald et al., 2013). This is particularly true for more symptomatic veterans, who (in contrast to control participants) showed no association between self-reported combat exposure and perceived threat (Figure 1c). In these more symptomatic veterans, perceived threat bias showed numerically higher correlations with anxiety, depression, and worry scores than perceived threat alone. This was due to paradoxical *inverse* correlations between these symptom measures and scores on the Combat Exposure Scale, which may have suppressed correlations between symptoms and perceived threat. We also found that perceived threat bias showed a specific association with avoidance/numbing symptoms, whereas perceived threat alone was correlated with total PTSD symptoms and scores on each subscale. This does not mean that perceived threat bias is a “better” measure of trauma-related pathology than perceived threat or combat exposure alone, but rather that it may tap into a unique facet of symptomatology that is not captured by investigating these sub-components in isolation.

The novel neurobiological contribution of this study was an inverse relationship between perceived threat while deployed and hippocampal volume (with no such relationships for amygdalar volume). It is unclear from these results whether it is absolute levels of perceived threat or perceived threat bias that is most relevant for hippocampal integrity. ROI analyses showed a significant relationship between perceived threat bias and hippocampal volume and a non-significant relationship for perceived threat. However, exploratory voxelwise analyses revealed a significant inverse association for perceived threat, and only a trend for perceived threat bias. While analyses with symptom measures showed numerically larger correlations with perceived threat bias than perceived threat, additional research is needed to demonstrate that this novel measure provides incremental validity over existing self-report measures. Replication and extension in a larger sample would allow for adequately powered tests of whether perceived threat bias accounts for greater variance in symptom measures or neurobiological features than does perceived threat. Future work should also investigate the predictive validity of perceived threat bias – for example, whether it shows unique associations with behavioral or functional outcomes, or whether discrepancies in trauma exposure and perceived threat predict responses to treatment.

Recognizing that perceived threat bias integrates two distinct underlying measures, we offer two accounts of psychological processes that may be involved. Perceived threat bias may reflect biased threat appraisals *during deployment*. High correlations between perceived threat bias and self-reported anxiety, depression, and worry suggest a relationship with previously described cognitive or attentional biases in mood and anxiety disorders. For example, perceived threat bias may reflect attentional bias to threat, observed in laboratory studies of PTSD and anxiety disorders (Bar-Haim et al., 2007). These disorders are also associated with interpretation bias, or a tendency to interpret ambiguous information as threatening (Yoon and Zinbarg, 2008). Although interpretation and attentional biases may be adaptive in unpredictable and potentially dangerous deployment contexts, such biases are less adaptive in safe, non-combat settings (Wald et al., 2013).

Alternatively, different individuals may appraise events similarly at the time of exposure, but retrospectively recall or report on these experiences very differently (Coles and Heimberg, 2002). For example, elevated perceived threat bias may be in part driven by underreporting of combat experiences. The positive relationship with CAPS avoidance symptoms lends support to the possibility that avoidance of aversive combat memories leads to underreporting (consistent with this avoidance account, we observed somewhat paradoxical *inverse* associations between combat exposure scores and symptoms of anxiety and depression in more symptomatic veterans; Table S2). Conversely, individuals with *negative* perceived threat bias may retrospectively minimize perceptions of threat experienced during deployment. Importantly, memories for specific traumatic events and perceived threat are not indelible, but evolve as traumatic events become more distal, particularly for individuals with elevated PTSD symptoms (Engelhard et al., 2008; Southwick et al., 1997).

These evolving memories – which largely depend on the integrity of the hippocampus – make it impossible to discern the sequence of events resulting in hippocampal alterations observed years after combat. To adjudicate between the accounts laid out above (or other alternatives), it will be important to conduct longitudinal research on perceived threat. By investigating hippocampal function and structure in relation to behavioral indices of hypervigilance and threat avoidance before, during, and after combat exposure, the causal relationship between hippocampal integrity and contextually inappropriate, maladaptive behavioral responses could be illuminated (Wald et al., 2013). Assessing perceived threat *during* deployment (Lancaster et al., 2016) along with objective indices of threat exposure (based on official military records; Wald et al., 2013) would reveal whether perceived threat bias assessed after combat reflects retrospective biases in perceived threat or reported combat exposure.

This work would address the outstanding question of whether reduced hippocampal volume predisposes individuals to perceive events as more threatening, or whether subjective perceptions of threat during deployment *contribute* to hippocampal damage, perhaps via chronic alterations to HPA axis output. Animal research demonstrates that chronically elevated levels of glucocorticoids cause cellular damage to the hippocampus (Magariños et al., 1996) observable at the macroscopic level (Uno et al., 1994). Human neuroimaging studies have found that basal plasma cortisol levels are inversely correlated with hippocampal volume (Lupien et al., 1998; Starkman et al., 1992), and that chronic stress is associated with reduced hippocampal volume 20 years later (Gianaros et al., 2007). Structural alterations of the hippocampus – whether a predisposing risk factor or consequence of perceived stress – would have deleterious consequences for hippocampal-dependent processes such as appropriate threat contextualization (Hayes et al., 2011; Liberzon and Abelson, 2016) and pattern separation ability (Kheirbek et al., 2012). The inability to ground threatening stimuli or fear memories in appropriate contexts may contribute to excessive avoidance of people, places, or things that bear resemblance to trauma-related stimuli, consistent with the relationship between avoidance symptoms and smaller hippocampal volume.

Notably, we did not observe a relationship between overall PTSD symptoms and hippocampal volume, although our relatively small sample size precludes strong conclusions from these null results. Work from van Rooij and colleagues (2015) further suggests that there are important individual differences beyond PTSD symptom severity reflected in hippocampal volume. These authors found no baseline volumetric differences between combat-exposed controls and PTSD patients who remitted following subsequent treatment, but smaller hippocampal volume in treatment-resistant PTSD patients. The authors concluded that reduced hippocampal volume is a risk factor for persistent PTSD, consistent with a classic report of smaller hippocampal volume in healthy identical twins of Vietnam veterans with PTSD (Gilbertson et al., 2002). Inflated perceived threat – which we found to be associated with smaller hippocampal volume – may be one factor that contributes to the persistence of PTSD. This suggestion is consistent with the identification here of significant relationships between avoidance/numbing symptoms of PTSD and both perceived threat bias and hippocampal volume, and previous observations of the central role of avoidance in the persistence of fear memories (Foa and Kozak, 1986).

### Limitations and future directions

A major limitation of this work is that the perceived threat bias index was constructed from two retrospective self-report measures collected on average 5 years after deployment, each of which is subject to response and recall biases. In addition to retrospective reporting errors, individuals with equivalent combat exposure scores may have experienced objectively different amounts of combat trauma, as these events vary in their duration and severity. The presence of PTSD symptomatology may systematically influence retrospective reporting, leading to inflated recall of perceived threat and misreporting of combat events. Combat exposure scores in the PTSS group were positively correlated with re-experiencing symptoms, which could reflect heightened estimates of combat exposure due to flashbacks or nightmares. These factors underscore the need for longitudinal studies and the incorporation of military records (or at least concurrent self-report measures) to assess combat exposure.

Our relatively small sample size increases the chances that the observed effect sizes are overestimates of true effect sizes in the population (Button et al., 2013; Yarkoni, 2009). Additionally, future research is warranted in larger and more diverse samples to validate relationships between perceived threat/perceived threat bias and hippocampal structure, and to extend these results to women, veterans of other conflicts, and non-combat trauma.

Finally, perceived threat bias scores are not absolute and must be interpreted relative to the specific sample being investigated. Future research should investigate whether a similar bias metric can be developed that is invariant of the specific sample, accompanied by the establishment of normative population distributions, which would increase the utility of this measure in both research and clinical applications.

### Summary

In summary, we have described a new construct for research on combat trauma, perceived threat bias. Perceived threat bias was associated with greater PTSD symptoms, self-reported anxiety, depression, and worry, and smaller hippocampal volume. These results suggest that the hippocampus is an important neural substrate for individual differences in subjective appraisals of threat, which have previously been shown to influence the development of PTSD and other mood and anxiety disorders. Future research on this construct that utilizes longitudinal neurobiological, behavioral, and symptom-based measures will elucidate the temporal milieu of events that result in elevated threat appraisal following trauma exposure.

Shaded areas indicate 95% confidence intervals.

## Supplementary Materials

**Supplementary Methods**

**Supplementary Results**

## Supplementary Tables

- **Table S1.** Correlations between perceived threat bias and its subcomponents and PTSD symptoms.
- **Table S2.** Correlations between perceived threat bias and its subcomponents and symptoms of anxiety, depression, and worry
- **Table S3.** Subcortical structure volumes for the full sample and individual subgroups

## Supplementary Figures

- **Figure S1.** Correlations between perceived threat bias and symptoms of depression, anxiety, and worry
- **Figure S2.** Correlations between hippocampus volume and scores on the PTSD-Checklist – Military Version and individual symptom clusters
- **Figure S3.** Voxelwise brain morphometry correlates of perceived threat bias and its subcomponents within the hippocampus and amygdala

## Supplementary Methods

Voxelwise brain morphometry (VBM) analyses were conducted using FSL-VBM (Douaud et al., 2007; http://fsl.fmrib.ox.ac.uk/fsl/fslwiki/FSLVBM). Structural images were brain-extracted using BET and segmented into white and gray matter. Gray matter images were registered to the MNI-152 standard space template using non-linear registration (FNIRT), and the resulting images were averaged and flipped along the x-axis to create a left-right symmetric, study-specific gray matter template. All subject-space gray matter images were non-linearly registered to this study-specific template and “modulated” to correct for local expansion (or contraction) due to the non-linear component of the spatial transformation, by multiplying each voxel of the registered gray matter image by the Jacobian of the warp field. The modulated gray matter images were then smoothed with an isotropic Gaussian kernel with a sigma of 3mm.

Small-volume-corrected analyses were conducted within the anatomically constrained amygdala and hippocampus, which were defined by thresholding Harvard-Oxford amygdala and hippocampus ROIs bilaterally at a 25% probability threshold and then combining these into a single ROI (Desikan et al., 2006). Voxelwise general linear models were applied to spatially smoothed gray matter maps using as independent variables perceived threat bias (PTB) scores, each of the components of PTB (perceived threat from the Deployment Risk and Resilience Inventory [DRRI] and Combat Exposure Scale scores), and Clinician-Administered PTSD Scale (CAPS) scores. Resulting statistical maps were corrected for multiple comparisons at *p* < 0.05 using permutation-based non-parametric testing (FSL’s randomise). We also conducted a search for whole-brain morphometric changes associated with each of these variables of interest.

## Supplementary Results

Within the anatomically defined amygdala and hippocampus, there were no significant clusters in which PTB or CAPS scores were associated with local morphometric changes. However, at a threshold of *p* < 0.1 (corrected), increased PTB was associated with contraction of subjects’ gray matter images (or reduced local volume) within the left hippocampus, particularly in the posterior hippocampus (Figure S3a).

This relationship was more robust and survived correction for multiple comparisons (*p* < 0.05, corrected) for perceived threat, which was inversely related to gray matter volume across much of the left hippocampus (Figure S3b). In the contralateral hemisphere, increased Combat Exposure Scale scores were associated with *increased* local gray matter volume in the right posterior hippocampus at a threshold of *p* < 0.1, corrected (Figure S3c).

No whole-brain significant clusters were identified for voxelwise regressions involving PTB, perceived threat or the Combat Exposure Scale, or CAPS scores.

## Supplementary Tables

**Table S1:** Pearson correlation coefficients between perceived threat bias and its subcomponents – the Combat Exposure Scale and the Perceived Threat subscale of the Deployment Risk and Resilience Inventory – and symptoms of PTSD. Correlations are presented for the entire sample (*N*=54) and the posttraumatic stress symptoms (PTSS) subgroup (*N*=36).

|  | Combat Exposure Scale |  | DRRI: Perceived Threat |  | Perceived Threat Bias |  |
| --- | --- | --- | --- | --- | --- | --- |
| ENTIRE SAMPLE (N=52) | *r*(50) | *p* | *r*(50) | *p* | *r*(50) | *p* |
| CAPS total | 0.20 | 0.15 | 0.40 | 0.004 | 0.16 | 0.25 |
| CAPS B (Re-experiencing) | 0.34 | 0.01 | 0.29 | 0.04 | -0.04 | 0.78 |
| CAPS C (Numbing/Avoidance) | 0.04 | 0.76 | 0.43 | 0.001 | 0.33 | 0.02 |
| CAPS D (Hyperarousal) | 0.21 | 0.13 | 0.30 | 0.03 | 0.07 | 0.62 |
| PTSS SUBGROUP (N=36) | *r*(34) | *p* | *r*(34) | *p* | *r*(34) | *p* |
| CAPS total | 0.07 | 0.65 | 0.15 | 0.39 | 0.04 | 0.82 |
| CAPS B (Re-experiencing) | 0.33 | 0.05 | 0.03 | 0.88 | -0.23 | 0.18 |
| CAPS C (Numbing/Avoidance) | -0.18 | 0.31 | 0.29 | 0.08 | 0.32 | 0.06 |
| CAPS D (Hyperarousal) | 0.12 | 0.49 | -0.06 | 0.73 | -0.13 | 0.46 |

**Table S2:** Pearson correlation coefficients between perceived threat bias and its subcomponents – the Combat Exposure Scale and the Perceived Threat subscale of the Deployment Risk and Resilience Inventory – and symptoms of depression, anxiety, and worry. Correlations are presented for the entire sample (*N*=54) and the posttraumatic stress symptoms (PTSS) subgroup (*N*=36). Across the entire sample, symptom measures were unassociated with combat exposure scores, and symptom correlations were comparable for perceived threat and perceived threat bias. In contrast, within the PTSS group, symptom measures were *negatively* correlated with combat exposure scores, and symptom correlations were numerically larger for perceived threat bias than for perceived threat.

|  | Combat Exposure Scale |  | DRRI: Perceived Threat |  | Perceived Threat Bias |  |
| --- | --- | --- | --- | --- | --- | --- |
| ENTIRE SAMPLE (N=52) | *r*(50) | *p* | *r*(50) | *p* | *r*(50) | *p* |
| Beck Depression Inventory | -0.12 | 0.40 | 0.46 | < 0.001 | 0.48 | < 0.001 |
| Beck Anxiety Inventory | -0.07 | 0.60 | 0.46 | < 0.001 | 0.44 | 0.001 |
| Penn State Worry Questionnaire | -0.10 | 0.50 | 0.48 | < 0.001 | 0.49 | < 0.001 |
| PTSS SUBGROUP (N=36) | *r*(34) | *p* | *r*(34) | *p* | *r*(34) | *p* |
| Beck Depression Inventory | -0.42 | 0.001 | 0.33 | 0.05 | 0.52 | 0.001 |
| Beck Anxiety Inventory | -0.31 | 0.07 | 0.31 | 0.06 | 0.43 | 0.009 |
| Penn State Worry Questionnaire | -0.29 | 0.08 | 0.41 | 0.01 | 0.48 | 0.003 |

**Table S3:** Mean and standard deviation of the volume of left, right, and bilateral hippocampus and amygdala for the entire sample (*N* = 52) and different subgroups of participants: combat-exposed controls (*N* = 16). the posttraumatic stress symptoms (PTSS) group (*N* = 36), and a subset of participants in the PTSS group with a PTSD diagnosis (*N* = 17). Values are presented in units of mm^3^.

|  | Left Hemisphere |  | Right Hemisphere |  | Bilateral |  |
| --- | --- | --- | --- | --- | --- | --- |
| HIPPOCAMPAL VOLUME | Mean | SD | Mean | SD | Mean | SD |
| All participants (*N* = 52) | 4461 | 422 | 4451 | 366 | 8912 | 760 |
| Combat-exposed controls (*N* = 16) | 4517 | 404 | 4479 | 360 | 8996 | 746 |
| PTSS group (*N* = 36) | 4436 | 432 | 4438 | 373 | 8874 | 774 |
| PTSD diagnosis (*N* = 17) | 4412 | 462 | 4384 | 395 | 8795 | 819 |
| AMYGDALAR VOLUME | Mean | SD | Mean | SD | Mean | SD |
| All participants (*N* = 52) | 1732 | 209 | 1956 | 253 | 3688 | 406 |
| Combat-exposed controls (*N* = 16) | 1747 | 227 | 1941 | 243 | 3688 | 445 |
| PTSS group (*N* = 36) | 1726 | 204 | 1963 | 261 | 3689 | 395 |
| PTSD diagnosis (*N* = 17) | 1722 | 241 | 1989 | 305 | 3712 | 453 |

## Supplementary Figures

**Figure S1:**
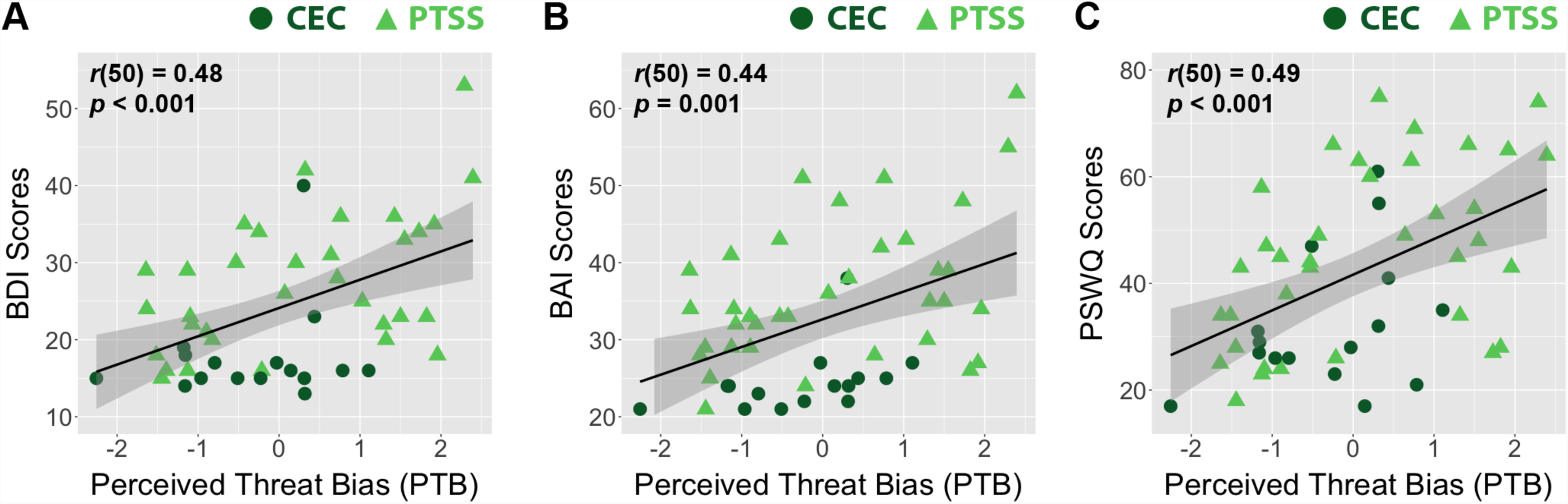
Perceived Threat Bias (PTB) scores were positively correlated with (A) self-reported symptoms of depression collected with the Beck Depression Inventory (BDI; entire sample *r*(50) = 0.48, *p* < 0.001; PTSS group *r*(34) = 0.52, *p* = 0.001), with (B) self-reported anxiety symptoms on the Beck Anxiety Inventory (BAI; entire sample *r*(50) = 0.44, *p* = 0.001; PTSS group *r*(34) = 0.43, *p* = 0.009), and with (C) self-reported worry characteristics from the Penn State Worry Questionnaire (PSWQ; entire sample *r*(50) = 0.49; *p* < 0.001; PTSS group *r*(34) = 0.48, *p* = 0.003). Shaded areas indicate 95% confidence intervals.

**Figure S2:**
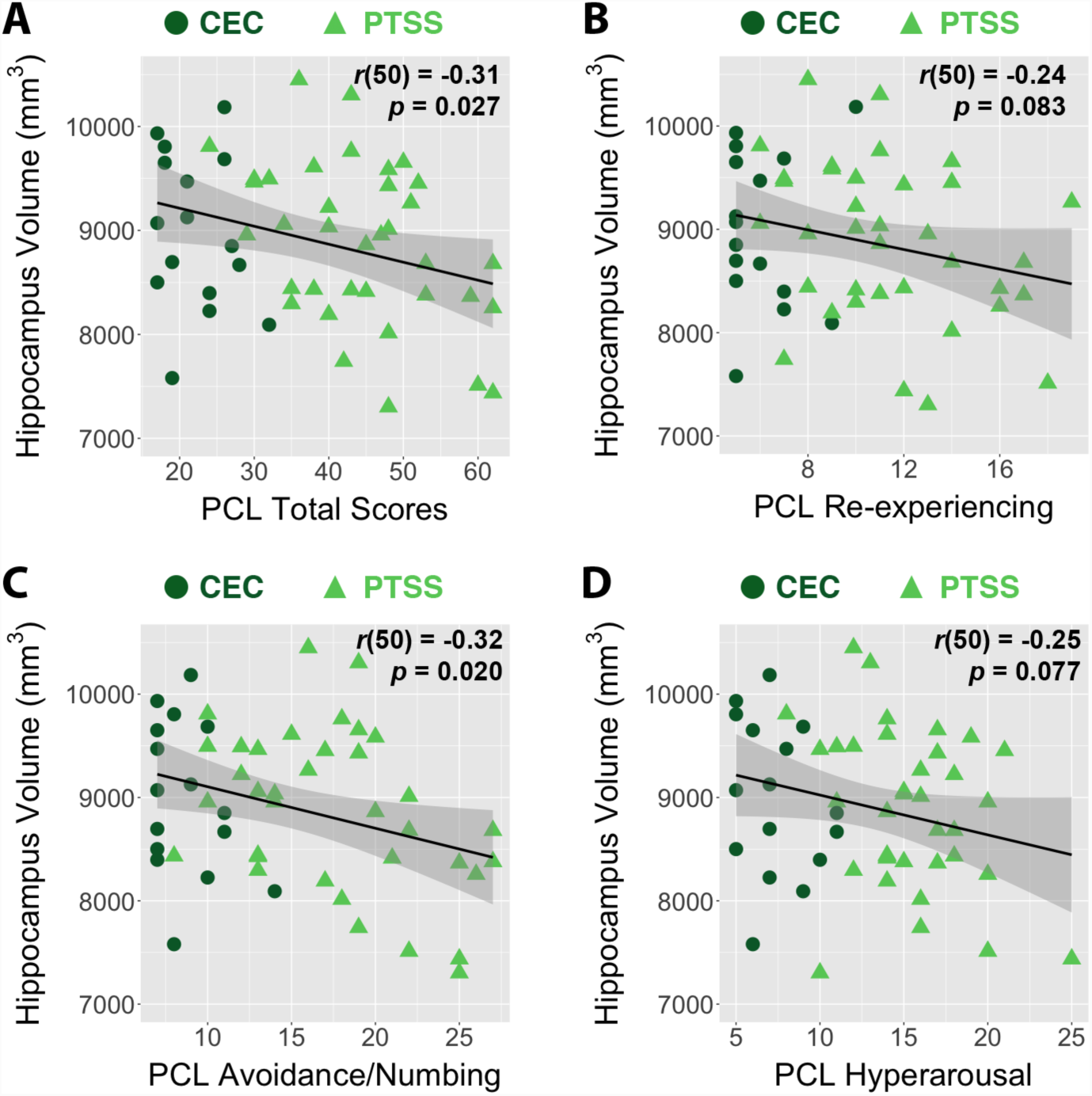
(A) Self-reported PTSD symptom severity on the PTSD Checklist-Military version (PCL), was significantly associated with smaller bilateral hippocampus volume across the entire sample (*r*(50) = -0.31, *p* = 0.027) and for subjects in the posttraumatic stress symptoms (PTSS) group alone (*r*(34) = -0.44, *p* = 0.0072). Negative relationships with hippocampus volume were observed for (B) re-experiencing symptoms (entire sample *r*(50) = -0.24; *p* = 0.083; PTSS *r*(34) = -0.33, *p* = 0.053), (C) avoidance/numbing symptoms (entire sample *r*(50) = -0.32, *p* = 0.020; PTSS *r*(34) = -0.41, *p* = 0.014), and (D) hyperarousal symptoms (entire sample *r*(50) = -0.25, *p* = 0.077; PTSS *r*(34) = -0.32, *p* = 0.060). Shaded areas indicate 95% confidence intervals.

**Figure S3:**
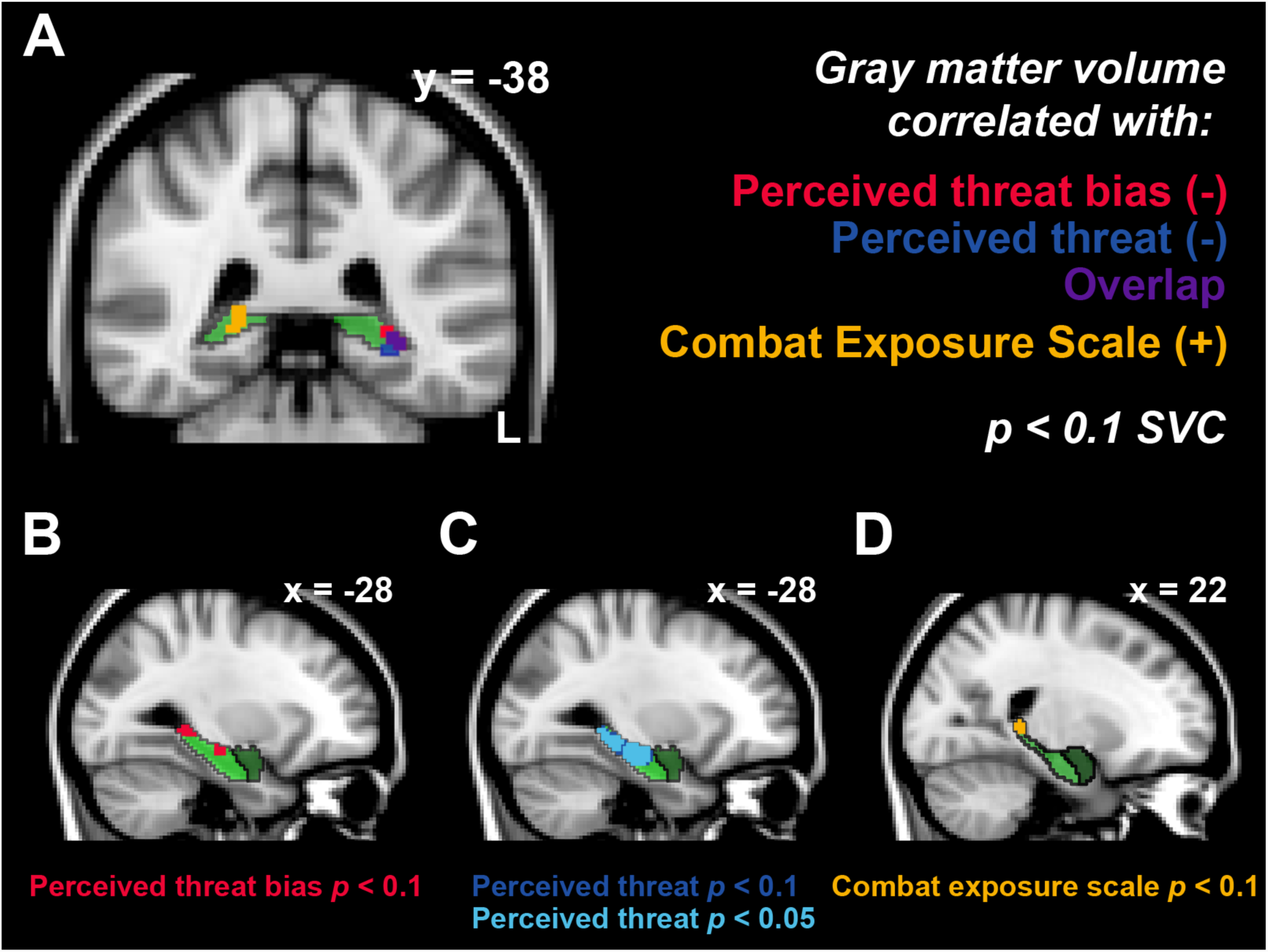
Regression of voxel-based morphometry values within the anatomically defined hippocampus (light green) and amygdala (dark green) on measures of interest. At a small-volume-corrected (SVC) threshold of *p* < 0.1, perceived threat bias was inversely related to local gray matter volume in the left posterior hippocampus (A, B). This relationship was driven by perceived threat scores (A, C), which showed a significant inverse relationship with gray matter volume (*p* < 0.05, SVC). Scores on the Combat Exposure Scale (A, D) were *positively* correlated with gray matter volume in the right posterior hippocampus (*p* < 0.1, SVC).

## Acknowledgements

The authors thank the participants for their military service and their involvement in this study, as well as the Wisconsin National Guard, the Madison VA Hospital, and other veterans’ community organizations for their assistance in recruitment. The authors also thank Andrew Fox and Regina Lapate for their insight in developing the perceived threat bias measure; Kate Rifken, Andrea Hayes, Emma Seppala, Kara Chung, Andy Hung, Michael Anderle, Lisa Angelos, Isa Dolski, Ron Fisher, and Nate Vack for their help with study planning, execution, and analysis; and Robin Goldman and Ryan Herringa for critical editorial feedback.

This work was financially supported by the Dana Foundation to JBN; the University of Wisconsin Institute for Clinical and Translational Research to Emma Seppala; the National Institute of Mental Health (NIMH) R01-MH043454 and T32-MH018931 to RJD; and a core grant to the Waisman Center from the National Institute of Child Health and Human Development (P30-HD003352). DWG was supported by a Graduate Research Fellowship from the National Science Foundation.

Portions of this work were previously presented at the 71^st^ Annual Scientific Convention of the Society of Biological Psychiatry, Atlanta, GA, May 12, 2016.

